# Defining endemism levels for biodiversity conservation: tree species in the Atlantic Forest hotspot

**DOI:** 10.1101/2020.02.08.939900

**Authors:** Renato A.F. Lima, Vinicius Castro Souza, Marinez Ferreira de Siqueira, Hans ter Steege

## Abstract

Endemic species are important for biodiversity conservation. Yet, quantifying endemism remains challenging because endemism concepts can be too strict (i.e., pure endemism) or too subjective (i.e., near endemism). We propose a data-driven approach to objectively estimate the proportion of records inside a given the target area (i.e., endemism level) that optimizes the separation of near-endemics from non-endemic species. We apply this approach to the Atlantic Forest tree flora using millions of herbarium records retrieved from multiple sources. We first report an updated checklist of 5044 species for the Atlantic Forest tree flora and then we compare how species-specific endemism levels obtained from herbarium data match species-specific endemism accepted by taxonomists. We show that an endemism level of 90% separates well pure and near-endemic from non-endemic species, which in the Atlantic Forest revealed an overall endemism ratio of 45% for its tree flora. We also found that the diversity of pure and near endemics and of endemics and overall species was congruent in space. Our results for the Atlantic Forest reinforce that pure and near endemic species can be combined to quantify regional endemism and therefore to set conservation priorities taking into account endemic species distribution. We provided general guidelines on how the proposed approach can be used to assess endemism levels of regional biotas in other parts of the world.

## 1. Introduction

One common practice in biodiversity conservation is to focus on species with high conservation value, such as species threatened with extinction (i.e., threatened species) or those exclusive to a given region or habitat (i.e., endemic species). Threatened and endemic species are important for conservation because they have a greater extinction risk than other species (Brooks et al., 2006; Myers et al., 2000; Peterson and Watson, 1998). In addition, the spatial patterns of total and endemic species richness can be congruent (Kier et al., 2009; Bonn et al., 2002; Storch et al., 2012), so prioritizing the protection of areas with high-levels of endemism could also safeguard the remaining biodiversity. However, there have been more efforts to delimit threatened species than endemic ones. Threatened species are grouped by clearly-defined categories, enclosed by objective criteria (IUCN, 2018), while species often are classified simply as being endemic or not.

There are proposals to divide endemics species based on spatial scale (e.g., narrow, regional and continental endemics), evolutionary history (e.g., neo and paleo endemics) or habitat specificity (e.g., edaphic endemics; Ferreira and Boldrini, 2011; Kruckeberg and Rabinowitz, 1985; Peterson and Watson, 1998). These proposals, however, implicitly assume that all individuals of a species are confined to a given region or habitat, also known as true or pure endemism (Tyler, 1996). If one record is found outside the target region, the species is to be (re)classified as non-endemic. Since pure endemism is rather strict, the term near-endemism has been used to describe species with few records outside the target region (Matthews et al., 1993; Carbutt and Edwards 2006; Platts et al., 2011; Noroozi et al., 2018). Near-endemics are the result of rare dispersal events, temporary establishment in different habitats or the existence small satellite populations (Matthews et al., 1993; Perera et al., 2011). It is important to emphasize that both types of endemism refer to species restricted to a specific area or habitat, which does not necessarily imply species with small extent of occurrence (<20,000 km^2^ *sensu* IUCN, 2018) or low local abundance (Rabinowitz, 1981).

The differentiation between pure and near endemics is challenging, because it may not be stable in time: near endemics can become pure endemics if habitat loss is higher outside than inside the target region (Carbutt and Edwards, 2006). Conversely, pure endemics may become near endemics with the accumulation of knowledge on their geographical distribution (Werneck et al., 2011). This is particularly true for geographically-restricted species, which often have scarce occurrence data. Furthermore, pure endemics may be classified as near endemics due to species misidentifications (Carbutt and Edwards, 2006) or by a questionable delimitation of the target region (Platts et al., 2011). In practice, conservation aims at protecting as many individuals as possible for a given species (IUCN, 2018). So, the differentiation between pure and near endemism may have little impact to plan conservation actions. Therefore, the question is: how to distinguish both groups of endemic species from non-endemic species? Defining pure endemism is straightforward, but separating near-endemics from non-endemic species can be quite subjective.

Here we propose a data-driven approach to objectively separate near-endemic from non-endemic species for conservation purposes. This approach can also be used to separate widespread species from occasional species, i.e., species frequent in other regions but sporadic in a given target region (Barlow et al., 2010). Therefore, its main goal is to classify species occurring inside a target region into pure-endemics, near-endemics, widespread and occasional species, which is done based on their ratio of occurrences inside the target region. As an example, we apply this approach to the Atlantic Forest, a global biodiversity hotspot with abundant knowledge on the taxonomy and distribution of its flora. We focus on the Atlantic Forest arborescent flora, a plant growth form which is well represented in biological collections (Daru et al., 2018). Using millions of carefully curated occurrences from over 500 collections around the world, we evaluate which ratio of occurrences inside the Atlantic Forest match species-specific endemism accepted by taxonomic experts. Finally, we illustrate the implications of the proposed approach to assess endemism ratio, to support on-the-ground conservation actions and to provide additional layer of information to existing tools of spatial prioritization.

## 2. Material and methods

### 2.1 An objective approach to delimit species endemism

Here, we formalize the six steps of the proposed approach to objectively classify species endemism levels, which can be applied in respect to any target region based on the distribution of species occurrence records (Figure 1).

**Figure 1.**
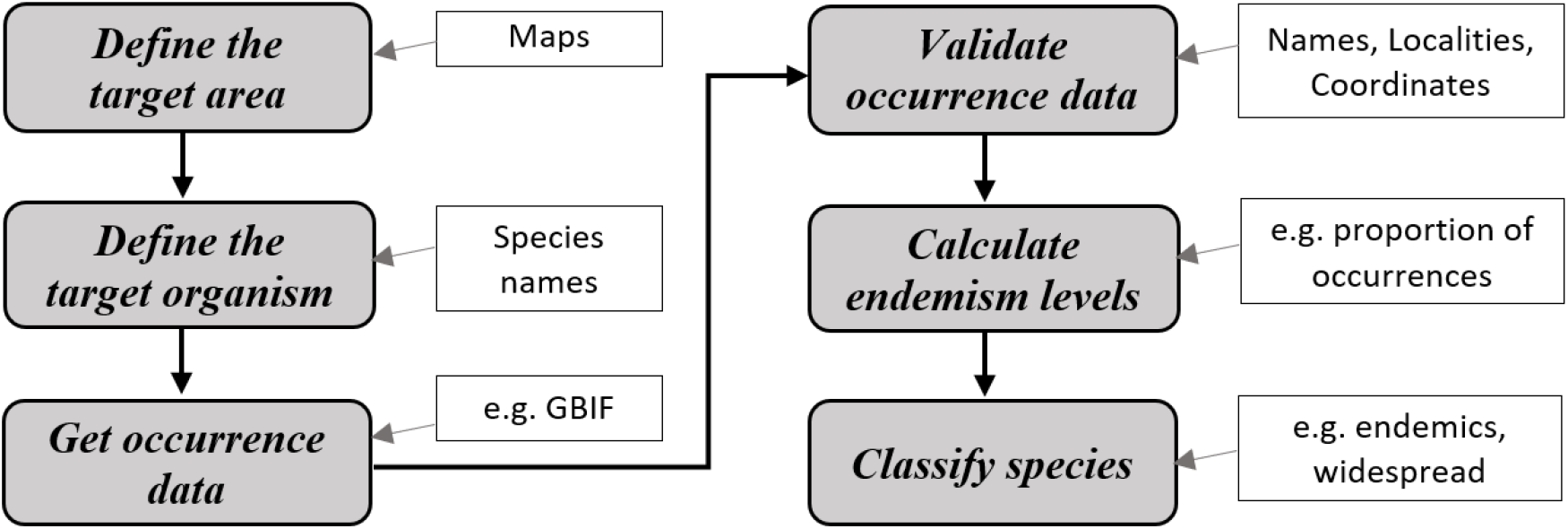
Flow chart showing the six steps of the proposed approach to classify species based on their endemism levels.

#### 2.1.1 Define the target area

Species endemism cannot be assessed without defining a geographical area. This area can be a region, domain or a habitat, but endemism is always relative and scale-dependent (Laffan and Crisp, 2003). Although many countries may want to produce their list of endemic species, it is recommended to use natural rather than political boundaries to define the geographical extent of the target area (Ferreira and Boldrini, 2011). If the target area is facing changes, such as an increasing loss of natural habitats, specifying the time window over which the target area is being considered may be relevant.

#### 2.1.2 Define the target organism(s)

Assessing the endemism of all living species is too time-consuming or data-limited for many taxa. Therefore, one needs to restrict the assessment to one or fewer taxa, that may be chosen according to their taxonomy (e.g., genus, family), or according to their life form, ecology (e.g., ecological guild), function (e.g., trophic levels) or conservation value (e.g., threatened species). Once the target organisms were defined, it is important to build a comprehensive list of names for all species occurring inside the target area. This list that should include synonyms and orthographical variants of the valid species names, to increase changes of occurrence data retrieval. If this a list of names is not available, a list of localities containing the target area (e.g., country names) can be used to generate a list of organisms potentially occurring inside the target area.

#### 2.1.3 Obtain species occurrence data

After defining the input list of names to search for species occurrences, it is necessary to define the data sources, which can be primary sources (e.g., personal field collections), secondary sources (e.g., biological collections, floras) or both. In the case of large databases of secondary sources (e.g., GBIF) and/or large number of taxa, the number of occurrences available may be large (thousands to millions). So, the use of automatized tools for data download and documentation may be needed (Chamberlain et al., 2020).

#### 2.1.4 Validate occurrence data

Particularly when using data from multiple secondary sources, it is important to validate the information accompanying the occurrences, such as the collector name and number, collection locality and geographical coordinates. In the case of two or more biological collections, the removal of duplicated specimens across collections is advised.

Depending on the characteristics of the data sources, one may need to remove spatial duplicates (e.g., records from the same localities) or spatial outliers (probable errors placed too far away from species core distributions). Another important validation step is to define the accepted confidence level of the taxonomic identification of each occurrence (e.g., use only identifications performed by taxonomists).

#### 2.1.5 Calculate species endemism levels

Next step simply is the count of the number of valid occurrences inside and outside the target area(s). This can be done by aggregating occurrences by locality names or by crossing a map of the target area with the geographical coordinates of the occurrences. The simplest endemism level metric possible is the number of valid occurrences inside the target area over the total number of occurrences retrieved for each taxon. If there is uncertainty on the delimitation of the boundaries of the target area (e.g., low-resolution map), the occurrences falling close to these boundaries may need a differential treatment to avoid biases on species classifications due to imprecise boundary delimitation (Platts et al., 2011).

#### 2.1.6 Classify species for conservation planning

The empirical levels of species endemism calculated in the previous step can be used as a metric of species endemicity in itself or as means to classify species into categories according to their degree of endemicity. For instance, if all records occur inside the target area the species can be classified as pure endemic and if the majority of the records occur outside, the species can be classified as occasional. One needs to assume (or estimate, see example below) thresholds of endemism level to separate near endemics and occasional from other species. Ideally, the classification should be compared to existing classifications of species endemism for validation.

### 2.2 A case study: tree species in the Atlantic Forest

We applied this approach to the arborescent flora of the Atlantic Forest biodiversity hotspot in eastern South America (see Supplementary Material for full details).

#### 2.2.1 Target area and organisms

The Atlantic Forest originally covered ca. 136 million hectares in three different countries, Argentina, Brazil and Paraguay (geographical range: 4‒34° S latitude, 35‒57° W longitude ‒ Figure S1a). Therefore, we searched for species occurrence data using a list of species names occurring in South America, compiled from different sources (Zuloaga et al., 2008; Oliveira-Filho, 2010; Grandtner and Chevrette 2013; Lima et al., 2015; Zappi et al., 2015; ter Steege et al., 2016). Here we considered only a part of the Atlantic Forest biota, the arborescent species, hereafter referred simply as trees. We considered tree species occurrences in all Atlantic Forest types, which include evergreen, semi-deciduous, deciduous, mixed temperate (locally known as *Araucaria* forests), white-sand (‘Restingas’ and ‘Mussunungas’), alluvial, cloud and swamp forests, as well as in rocky field and inselberg vegetation. Arborescent species are relatively well represented in herbaria (Daru et al., 2018) and they are defined here as species with free-standing stems exceeding 5 cm of diameter at breast height (1.3 m) or 4 m in total height, including arborescent palms, cactus, tree ferns, and woody bamboos. Moreover, some tall shrubs and treelets are included here under the term trees. We carefully inspected the input list of names to avoid the inclusion of exotic and non-arborescent species.

#### 2.2.2 Retrieval and validation of occurrence data

The list of South American tree names was used to download occurrence data from multiple secondary sources, namely *species*Link (www.splink.org.br), JABOT (http://jabot.jbrj.gov.br, Silva et al., 2017), ‘Portal de Datos de Biodiversidad Argentina’ (https://datos.sndb.mincyt.gob.ar) and the Global Biodiversity Information Facility (GBIF.org, 2019). We excluded all occurrences described in the specimen notes as being cultivated or exotic. We checked names for typos, orthographical variants and synonyms in the Brazilian Flora 2020 (BF-2020) project (Filardi et al., 2018; Zappi et al., 2015). Decisions for unresolved names were made by consulting Tropicos (www.tropicos.org) or the World Checklist of Selected Plant Families (http://wcsp.science.kew.org).

There was much variation of the notation across herbaria, on the locality details provided and on the precision of the geographical coordinates among the millions of records retrieved (Appendix A). Therefore, we conducted a detailed data cleaning and validation procedure (see Supplementary Material for details). We standardized the notation of different fields (e.g., locality description, collector and identifier names, collection and identification dates), which were then used to (i) search for duplicate specimens among herbaria; (ii) validate the geographical coordinates at country, state and/or county levels and (iii) to assess the confidence level of the identification of each specimen (i.e., ‘validated’ and ‘probably validated’ identifications -Appendix B).

Moreover, (iv) we cross-validated information of duplicate specimens across herbaria to obtain missing or more precise coordinates and/or valid specimen identifications. Finally, (iv) we removed specimens too distant from their core distributions (i.e., spatial outliers), which are often related to specimens collected from cultivated individuals but that are not declared so by the collectors.

#### 2.2.3 Calculating species endemism levels

We calculated an empirical level of endemism based on the position of records for each species in respect to the Atlantic Forest limits (Olson and Dinerstein, 2002; IBGE, 2012). Each record was assigned as being inside, outside or in the transition of the Atlantic Forest to other domains (see details in Figure S1b). Records in the transition were those falling inside the Atlantic Forest limits, but in counties with less than 90% of its area inside the Atlantic Forest or vice-versa. Because of the variable precision of the specimen’s coordinates and of the uncertainty of the boundary delimitation at the scale of our target area map (1:5,000,000), records in the transition received half the weight other records to calculated species endemism levels:

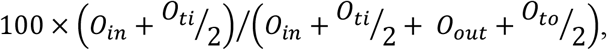

where, *O*_in_, *O*_ti_, *O*_out_ and *O*_to_ are the number of specimens inside, inside in the transition, outside and outside in the transition to the Atlantic Forest, respectively. This endemism level is actually a weighted proportion of occurrences inside the Atlantic Forest by the total of valid occurrences found, varying from 0 (no occurrences) to 100% (all occurrences inside the Atlantic Forest).

We then obtained the endemism classification derived from the expertise of taxonomists working on the BF-2020 project (Filardi et al., 2018), the best reference currently available for the Atlantic Forest flora. Each species was classified as ‘endemic’ if the BF-2020 field ‘phytogeographic domain’ contained only the term ‘Atlantic Rainforest’ (equivalent to what we refer here as Atlantic Forest with all of its forest types). Correspondingly, a species was classified as ‘occasional’ if this field did not include this term. Species with no information on the ‘phytogeographic domain’ were omitted from this analysis.

The comparison between the empirical classification of species endemism and the reference BF-2020 classification was based on thresholds values varying from 0 to 100%, in intervals of 1% (i.e., 0, 1, …, 99, 100%). If a given species had an observed endemism level equal or higher than a given threshold, it was classified as ‘endemic’. For each threshold value, we calculated the number of mismatches between the two classifications (i.e., species classified as ‘endemic’ in the BF-2020 and ‘not endemic’ from the observed endemism level or vice-versa). The same procedure was used to calculate the number of mismatches for occasional species. We then plotted the number of mismatches against all thresholds and estimated the optimum threshold that minimizes the number of mismatches between classifications. Optimum thresholds were estimated using piecewise regression, allowing up to five segments (i.e., four breaking points). Thus, we provided the breaking point of each curve (and its 95% confidence interval). We compared the results using only taxonomically ‘validated’ and using both taxonomically ‘validated’ and ‘probably validated’ records.

#### 2.2.4 Species classification and implications for conservation planning

We used the optimum threshold values obtained above to classify species into pure endemics, near endemics, widespread and occasional species. Because endemic species are not necessarily narrowly distributed and occasional species may be frequent elsewhere, this terminology tried to reflect broad patterns of species occurrence in respect to the target region (pure and near endemics: all or nearly all occurrences within the target region; widespread: species with many occurrences both within and outside the target area; occasional: species with most occurrences outside the target area). We then used this classification to delimit the centers of diversity for each group of species (Laffan and Crisp, 2003). In order to do so, we plotted the valid occurrences of each group of species against a 50×50 km grid covering the Atlantic Forest and surrounding domains. Next, we obtained different diversity metrics for each group of species per grid cell. We selected two metrics with best performance to describe our data (Figures S2 and S3): corrected weighted endemism (WE) and rarefied/extrapolated richness (SRE). The WE is the species richness weighted by the inverse of the number of cells where the species is present, divided by cell richness (Crisp et al., 2001). The SRE is the rarefied/extrapolated richness (depending on the observed number of occurrences per cell) for a common number of 100 occurrences, calculated based on the species frequencies per cell (Chao et al., 2014). We also obtained the sample coverage estimate (Chao and Jost, 2012), used here as a proxy of sample completeness. We evaluated the relationship of the diversity of endemic and occasional species with overall species diversity using spatial regression models (i.e., linear regression with spatially correlated errors - Pinheiro and Bates, 2000). Centers of diversity were delimited using ordinary kriging and only the grid cells meeting some minimum criteria of sampling coverage (see Supplementary Material). We used the 80% quantile of predicted distributions to delimit the centers of endemism.

## 3. Results

The search for occurrence records based on this input list of tree names resulted in a total of 3.11 million records from 543 collections (Appendix A). After the removal of duplicates, spatial outliers and the geographical and taxonomic validation, we retained 593,920 valid records (disregarding records with ‘probably validated’ taxonomy) for the classification of species endemism. We found 252,911 valid records being collected inside the Atlantic Forest limits, which contained a total of 5044 arborescent species (4054 species excluding tall shrubs; Appendix C). If we consider the valid occurrences in the transitions of the Atlantic Forest to other domains, we could add 294 species as probably occurring in the Atlantic Forest (Appendix D). Another 3158 names were retrieved but were finally excluded from the list for different reasons (e.g., synonyms, typos, orthographical variants, species not occurring naturally in the Atlantic Forest, etc.; Appendix E).

Based on the valid records retrieved for the Atlantic Forest, we found evidence of pure endemism (i.e., endemism level= 100%) for 1547 tree species (31%; Appendix F). We found that 90.2% of records inside the Atlantic Forest (95% Confidence Interval, CI: 89.3‒91.2%) was the threshold of endemism level that best matched the endemism currently accepted by taxonomy experts (Figure 2a). The curve of mismatches between the observed and reference classifications decreases until it reaches a minimum and then it increases again, meaning that more or less restrictive thresholds lead to an increase the number of mismatches. The 90.2% threshold in the Atlantic Forest added 733 near endemic species (15%). Together, pure and near endemics lead to an overall endemism ratio of 45.2% for the Atlantic Forest arborescent flora (Figure 2b) and 1.01 endemic arborescent species per 100 km^2^ of remaining forest (i.e., 2261.2 km^2^; Fundación Vida Silvestre Argentina and WWF, 2017). Conversely, we found that 8.7% (95% CI: 8.2‒9.3%) was the best threshold for separating occasional from widespread species occurring in the Atlantic Forest (Figure 2a), leading to a total of 639 occasional species (13%). The remaining 42% of the species were classified as widespread (Appendix F). Results using only occurrences with taxonomy flagged as ‘validated’ were similar (pure endemism: 32%; near endemism: 15%; occasional species: 14%, widespread species: 39% -Figure S4, Appendix F).

**Figure 2.**
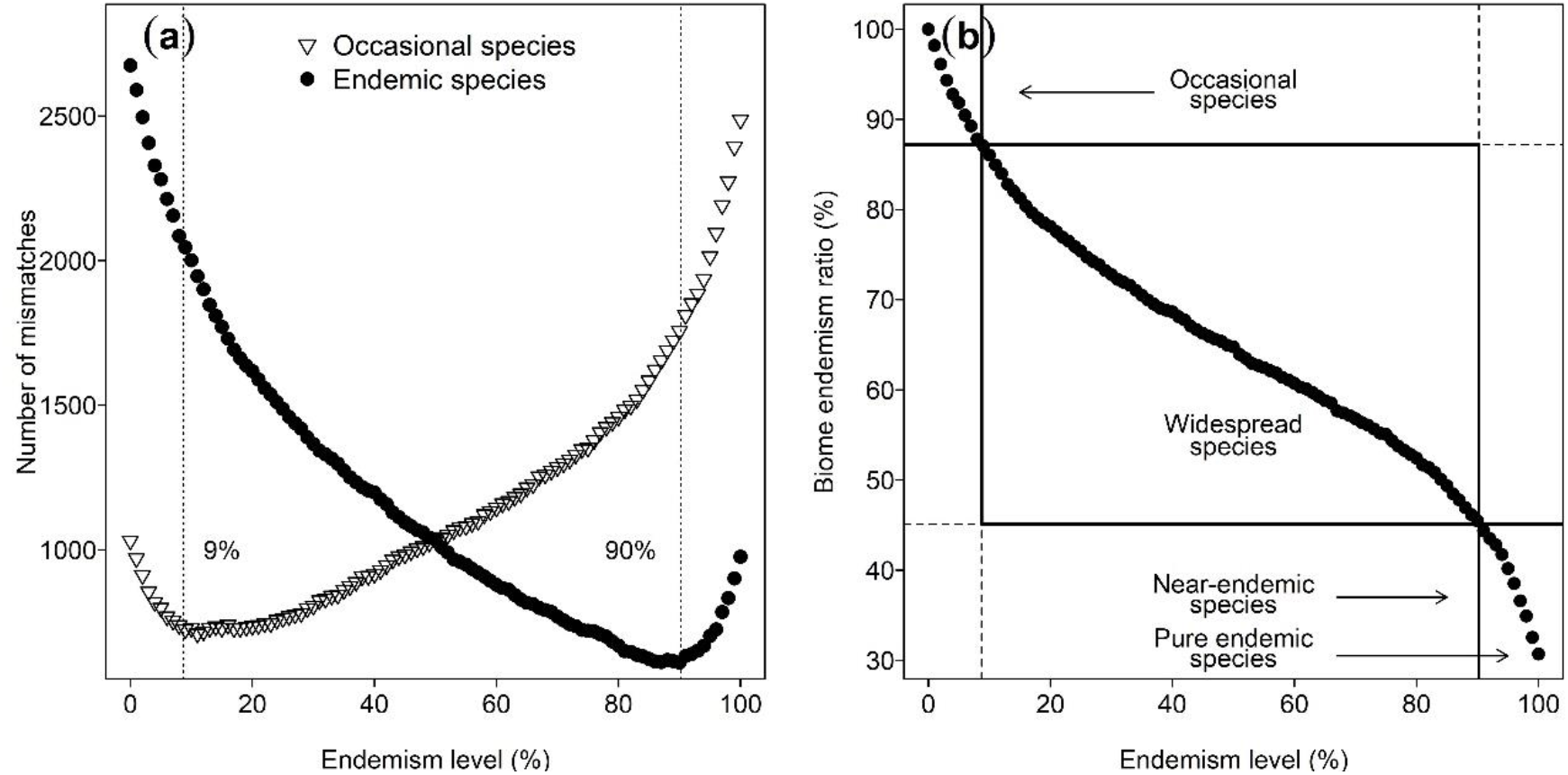
Defining near endemic and occasional tree species using herbarium records for the Atlantic Forest biodiversity hotspot. For both endemic (black circles) and occasional species (triangles), we present (a) the optimum endemism levels (vertical dashed lines) estimated from the distribution of mismatches between the empirical and the Brazilian Flora 2020 classifications and (b) the overall endemism ratio of the Atlantic Forest in intervals of 1% (*x*-axis in both panels).

The diversity of endemic species was strongly correlated with the overall species diversity in the Atlantic Forest (Figure 3). There was also a strong and positive correlation between the number of pure and near endemic species (Figure S5), meaning that the centers of diversity of pure and near endemics are highly congruent in space. The diversity of pure endemics was higher in the rainforests along the coast (Figure 4), corresponding to the rainforests of the Serra do Mar and Bahia Coastal Forests ecoregions (Olson and Dinerstein, 2002). The inclusion of near endemics expanded the diversity of endemic species towards more inland parts of the Atlantic Forest, but spatial patterns remained quite similar (Figure 4 and Figures S6-S8). This expansion was more conspicuous in the colder Araucaria forests in the southern Atlantic Forest, but not to the point of including these forests as centers of diversity (i.e., areas with the 80% higher values). On the other hand, occasional species were really rare in the Araucaria forests. Most of the distribution of occasional species was concentrated in the Brazilian Cerrado, but also in the Amazon and slightly less in the Caatinga domain. General patterns were fairly similar when using other diversity measures (Figures S6-S8).

**Figure 3.**
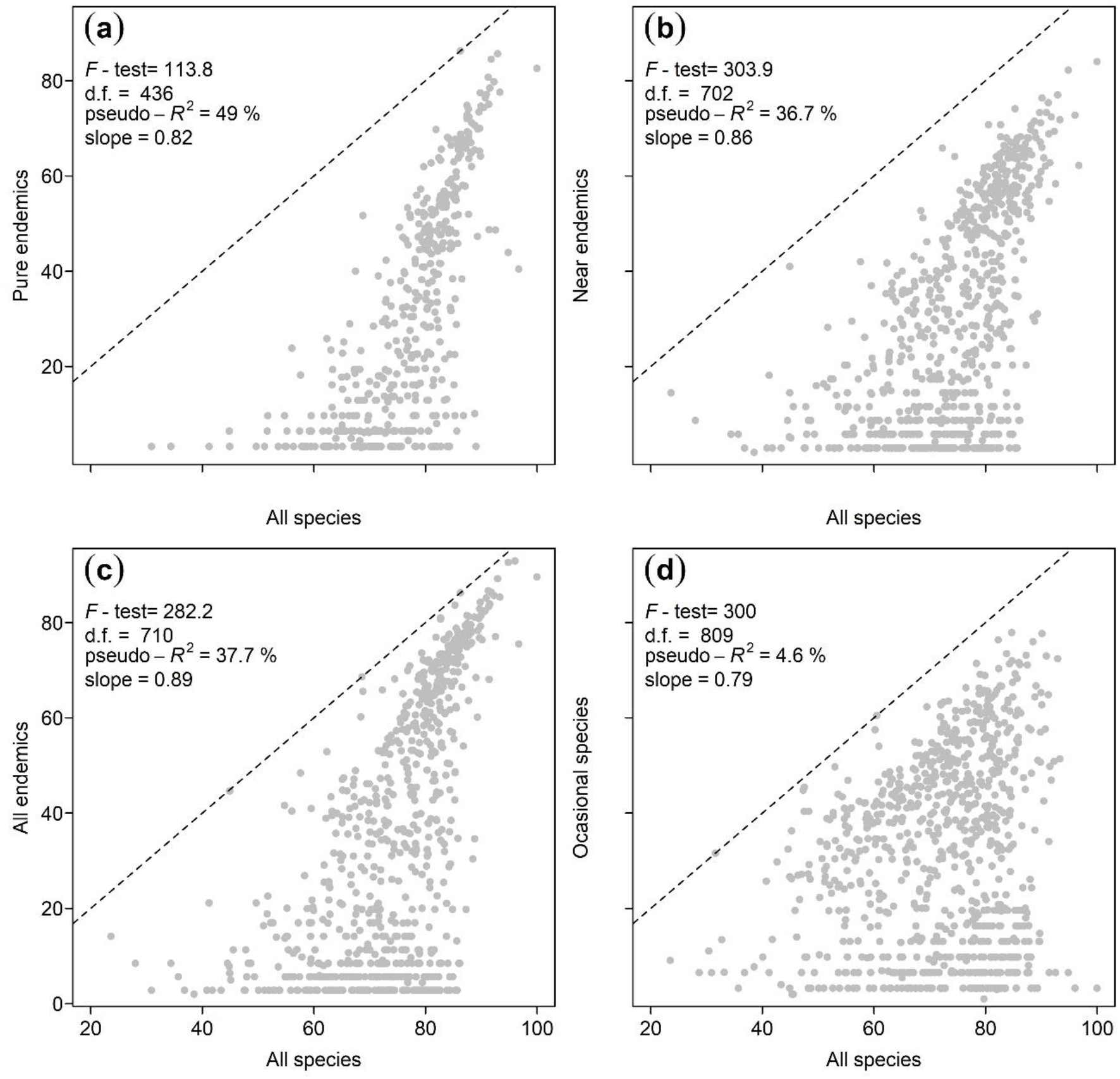
Relationship between the number of rarefied/extrapolated richness per 50×50 km grid cell and the same diversity metric obtained for (a) pure endemics, (b) near endemics, (c) all endemics (pure + near endemics) and (d) occasional species. For each group of species, we present the summary statistics of each spatial regression model (top left; d.f.= degrees of freedom), including the predicted slope of the regression prediction. The spatial regression analysis was performed only for grid cells meeting some minimum criteria of sampling coverage (see Supplementary Methods). The dashed line represents the 1:1 line. All *p*-values are below 0.001.

**Figure 4.**
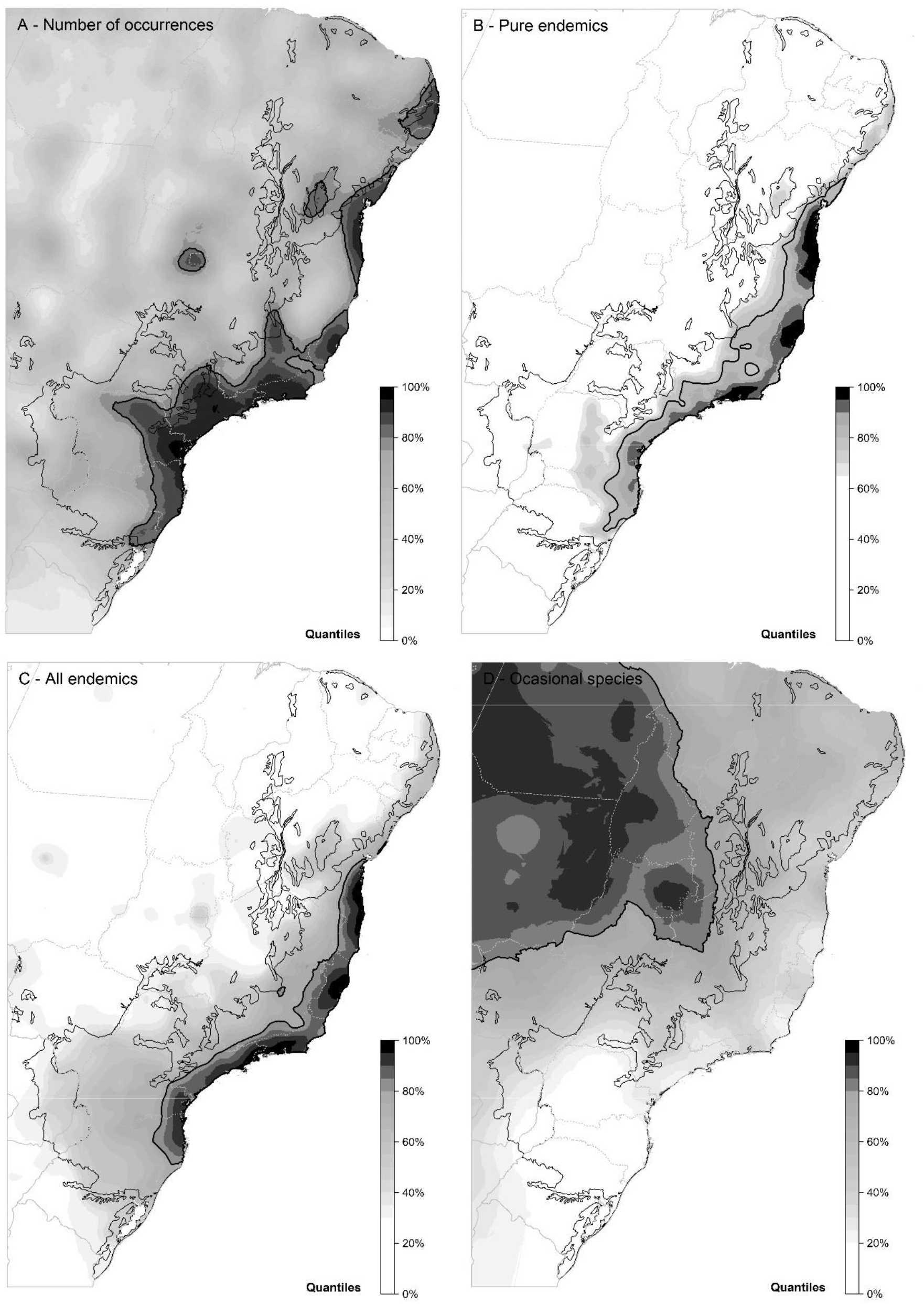
The spatial distribution of (A) the number of occurrences retrieved for the species occurring in the Atlantic Forest, and the centers of diversity of (B) pure endemics, (C) all endemics (pure + near) and (D) occasional species. Maps were produced using ordinary kriging based on rarefied/extrapolated species richness obtained for a common number of 100 records per grid cell. The color scale represents the 5% quantiles of the metrics distribution, from 0-5% (white) to 95-100% (black). Bold black lines are the area containing the 80% higher richness values. The black line marks the limits of the Atlantic Forest, while the solid and dashed grey lines mark the limits of South American countries and of the Brazilian states, respectively.

## 4. Discussion

### 4.1 Describing species endemism

Near endemism has been used to assess endemism levels of regional floras and faunas. However, such assessments often use loose (Carbutt and Edwards, 2006; Platts et al., 2011) or arbitrary definitions (Perera et al., 2011; Noroozi et al., 2018) of near endemics. Here, we propose and apply an objective approach to find that 90% of the occurrences inside a target region can be used to tell apart endemic species from non-endemic species, a result supported by endemism classifications performed by taxonomic experts. This 90% limit has one important implication: the average endemism concept adopted by taxonomic experts implicitly includes the concept of near endemism, at least for the Atlantic Forest. Indeed, the overall endemism ratio found here for pure and near endemics combined (45%) is within the range of 40-50% endemism level previously reported for the flora of this biodiversity hotspot (Myers et al., 2000; Stehmann et al., 2009; Zappi et al., 2015). Thus, we propose that pure and near endemics can be used together to objectively delimit endemism or as two categories of endemism, similarly to what already exists for the categories of species threat (IUCN, 2018). Moreover, conservation funding is not always aligned with the degree of species endemism (Martín-López et al., 2009), despite the civic and scientific awareness of the role of endemics for prioritizing conservation (Myers et al., 2000; Brooks et al., 2006; Meuser et al., 2009; see Scarano, 2009 for a different point of view). Thus, we hope that the quantitative description of endemism proposed here can help to bridge the scarcity of conservation actions using information on species endemicity.

The threshold of 90% found here was also used to assess plant endemism in the Mediterranean Basin biodiversity hotspot (Médail and Baumel, 2018), suggesting that this threshold could be used in the assessment of plant endemism of other species-rich regions. However, we did not find similar assessments in the literature to confirm this suggestion. Thus, although our approach to delimit species endemism is objective and more comprehensive than pure endemism, similar assessments in other parts of the world and for other groups of species are still needed. We provide a workflow to perform such assessments, which would require (*i*) a list of species names, (*ii*) available sources of occurrence data, (*iii*) a data cleaning/validation pipeline, (*iv*) a digitized map of the study area, and (*v*) a classification of endemism based on taxonomists expertise. Online occurrence data sources (e.g., GBIF) and tools to download data (e.g., Chamberlain et al., 2020) and validate their geographical coordinates (e.g., Zizka et al., 2019) are becoming increasingly available. Here, we propose a simple but efficient way to validate the taxonomic determinations of specimens (see Supplementary Material). The bottleneck for applying this approach remains on the availability of regional lists of species names and on the quantity and accessibility of data from local collections (Boakes et al., 2010). These constraints may become more restrictive in species-rich and less economically developed regions. The Atlantic Forest, used here as a testing ground to our proposed approach, combines one of the largest number of species occurrences available for the tropics (see details below), with one of the most completed national floras (i.e., expert endemism information available ‒ Brazilian Flora project) and herbaria networks (e.g., *species*Link, JABOT).

### 4.2 Implications for conservation

The application of our approach to the tree flora of the Atlantic Forest offers insights on how it can be used for supporting the conservation of local floras or faunas. The first insight is related to the total number of species reported to a given region. The Atlantic Forest is arguably the tropical forest with one of the largest botanical knowledge available, with ca. 680,000 unique specimens of tree species, or 42 specimens per 100 km^2^ ‒ average collection density in the Amazon forest is below 10 per 100 km^2^ (ter Steege et al., 2016). Nevertheless, we over 700 new valid occurrences of tree species for this biodiversity hotspot, an increase of 21% to the 3343 trees previously reported by the Brazilian Flora 2020 project (Zappi et al., 2015). About 47% of these new records were represented by occasional species, which correspond to 13% of the total richness of the Atlantic Forest tree flora. This result confirms that occasional species, despite of their infrequency, make an important contribution to overall biodiversity of regional biotas (Barlow et al., 2010; ter Steege et al., 2019). But more importantly, 53% of the new records correspond to widespread species and endemic species. An increase of 16% in the total richness was also observed for the Espírito Santo state flora compared to the reported in the Brazilian Flora (Dutra et al., 2015). The Brazilian Flora 2020 project is permanently being improved and is of utmost importance for the understanding of the Brazilian flora (Zappi et al., 2015; Filardi et al., 2018), the richest in the world (Ulloa et al., 2017). Here, we provide products that can be readily integrated into the Brazilian Flora project (e.g., more refined endemism filters), illustrating how data-driven approaches as the one proposed here can help to refine the knowledge of regional floras, even in regions with a great knowledge about its flora, promoting the accumulation of critical knowledge to support biodiversity conservation.

Another possible application of the approach is the detection of centers of endemic species diversity. In the Atlantic Forest example provided here, the centers detected were congruent with previous proposals, which suggested areas of high endemism in the moist and rain forests between the Brazilian states of São Paulo and Rio de Janeiro and between Espírito Santo and Bahia states (Thomas et al., 1998; Murray-Smith et al., 2009). However, our results provided evidence that the coastal lowland forests in the states of Paraná and Santa Catarina (PR-SC) should also be included as important centers of tree endemism for the Atlantic Forest. In accordance to Murray-Smith et al. (2009), we found no strong support for the existence of an area of endemism along the coastal and ‘*brejo de altitude*’ forests in Paraíba, Pernambuco and Alagoas states (Thomas et al., 1998), at least not at the spatial scale used here (50×50 km). The Atlantic Forests of northeast Brazil are closer or are surrounded by seasonally dry vegetation (i.e., *Caatinga*) and they share many floristic elements with Amazon forests (Santos et al., 2007), which could lead to the lower endemism levels found for the species occurring in this part of the Atlantic Forest.

The provision of lists of species along with their degree of endemicity can support the selection of species for conservation projects (Martín-López et al., 2009; Meuser et al., 2009). These projects could be related to on-the-ground actions targeting individual species (e.g., Martins, 2014 or www.saveourspecies.org) or to restoration plans aiming at the maximization of biodiversity conservation outcomes while restoring ecosystem services (Brancalion et al., 2018). Moreover, since range-restricted endemics are probably also threatened, existing initiatives such as the Brazilian Alliance for Extinction Zero (www.biodiversitas.org.br/baze) could incorporate the information on degree of endemicity in their species selection methods. It is important to emphasize that not only the degree of endemicity should be taken into account in the selection of species for conservation projects. Widespread species may play important functional roles in natural ecosystems, so they should be included in conservation projects as well (Scarano, 2009).

The delimitation of centers of endemic diversity also has direct implications for conservation planning. For instance, they can assist the identification of Important Plant Areas (IPA), provided by the Target 5 of the Global Strategy for Plant Conservation (www.cbd.int/gspc), or of Key Biodiversity Areas (KBA - www.keybiodiversityareas.org). Although the delimitation of IPAs and KBAs predicts the use of endemic species, their definition is mainly based on the presence of threatened species. Also, IPAs are highly concentrated non-tropical regions of the northern hemisphere (www.plantlifeipa.org). Our data driven approach, based on careful data curation, proved to be efficient to identify areas of high endemicity in one of the richest tropical floras of the world and could be used to expand the IPA and KBA programs. In the specific case of the Atlantic Forest, which has less than 20% of its original forest cover, conservation actions are urgently needed. When combined with other layers of information (e.g., socio-economic), maps of endemic species diversity can be used as an additional layer of biodiversity information in existing tools of spatial prioritization (e.g., Brancalion et al., 2019; Strassburg et al., 2019), aiming to pinpoint remaining natural areas that should be protected or degraded lands that could be prioritized in restoration actions. This suggestion is reinforced by the spatial congruence found between the diversity of endemic and non-endemic tree species, meaning that conservation of areas with high-levels of endemism could also safeguard a great deal of the remaining Atlantic Forest tree flora (Kier et al., 2009; Bonn et al., 2002). Thus, considering that defining threatened and endemic species have the same constraints related to data availability and to the time and spatial scale considered (Ferreira and Boldrini, 2011), the detection of endemics is more straightforward than threatened species, which could speed up the decision-making process for conservation in rich tropical biotas around the world.

## Data Availability

All data providers and their citations are given in Appendix A. GBIF data used in the analysis is also provided in the references.

## CRediT authorship contribution statement

Renato A. F. de Lima: Conceptualization, Methodology, Formal analysis, Data curation, Funding acquisition, Writing - original draft. Vinicius C. Souza: Validation, Data curation, Writing - review & editing. Marinez F. Siqueira: Methodology, Writing - review & editing. Hans ter Steege: Methodology, Funding acquisition, Writing - review & editing.

## Acknowledgments

We thank Sidnei Souza and Renato Giovanni for their help with data compilation from speciesLink network. We also thank Lucie Zinger for helping with GBIF data management and for her suggestions on this manuscript. This study was supported by the European Union’s Horizon 2020 research and innovation program under the Marie Skłodowska-Curie grant agreement No 795114.

## Appendices

## Appendix A

List of collections and data providers used for data compilation.

The numbers of records retrieved per collection correspond to overall sum of records before data validation, thus including both valid and invalid records.

## Appendix B

List of names of taxonomists per family used for taxonomical validation. The ‘tdwg.name’ represents the taxonomist name following the standard notation of the Biodiversity Information Standards (https://www.tdwg.org), which includes different variants of notation found for the same taxonomist name.

## Appendix C

Updated, taxonomically vetted checklist of the Atlantic Forest tree flora. For each name included in the checklist we provide the life form, the status of the name in respect to the Brazilian Flora 2020 project, the number of records found inside the Atlantic Forest (both ‘validated’ and ‘probably validated’ taxonomy) and a list of up to 30 vouchers (only specimens with ‘validated’ taxonomy), giving priority to type specimens. We also indicate which species were regarded as being taxa of low taxonomic complexity (TBC) or taxa commonly cultivated outside its original range.

## Appendix D

List of species with probable occurrence in the Atlantic Forest.

We present all names with valid records found only in the transition of the Atlantic Forest to other domains and those names cited in the Brazilian Flora 2020 project as being an Atlantic Forest species, but for which we did not find any valid records. Again, we present for each name the life form, the number of records found and a list of up to 30 vouchers.

## Appendix E

List of names excluded from the final Atlantic Forest checklist.

For each name on the list we provide the life form and the reason why the name was excluded. For synonyms, orthographical variants, common typos we also provide the corresponding valid name used in this study.

## Appendix F

Endemism levels for the Atlantic Forest tree flora and the corresponding classification into pure endemic, near endemic, widespread and occasional species.

For each species name, we provide the number of valid records outside the Atlantic Forest, outside but in the transition to the Atlantic Forest, inside the Atlantic Forest but in the transition to other domains, and inside the Atlantic Forest. We present the endemism levels and species classifications using only records with validated taxonomy and using records with validated and probably validated taxonomy. Finally, we present the endemism classification currently accepted in the Brazilian Flora 2020 in respect to the Atlantic Forest.

## Appendix G

Shapefiles delimiting the centers of the endemic and occasional species diversity in the Atlantic Forest for pure endemics, near endemics, pure + near endemics and occasional species.

Each shapefile contains the isoclines corresponding to the 75%, 80%, 85%, 90% and 95% quantiles of the distribution of rarefied/extrapolated richness for 100 specimens, predicted using ordinary kriging.

