## Supplementary Information for "Defining endemism levels for biodiversity conservation: tree species in the Atlantic Forest hotspot"

**This PDF file includes:**

- **Supplementary Methods and References**
- **Supplementary Figures**

### SUPPLEMENTARY METHODS

#### The input list of tree names for South America

Searching for occurrence records depends on the input list of species names. Because the Atlantic Forest covers Argentina, Brazil and Paraguay (Figure S1), we started with a list of names of native trees of South America compiled by Grandtner and Chevrette (2013). We crossed this preliminary list of names with other sources to retrieve missing names: the Brazilian Flora 2020 (version 393.158) downloaded in June 2018 (http://reflora.jbrj.gov.br, Ranzato Filardi et al., 2018); and the Southern Cone Flora projects downloaded in July 2018 (http://www.darwin.edu.ar, Zuloaga et al., 2008). We also obtained missing names from other projects working with tree composition and diversity in the Neotropics, namely the Amazon Tree Diversity Network (ter Steege et al., 2016), NeoTropTree (Oliveira-Filho, 2010), and TreeCo (Lima et al., 2015). We added to this list some common typos (*e.g*., *Copaifera langsdorfii* for *Copaifera langsdorffii*), to avoid the exclusion of occurrences with misspelled names. The final input list had 66,895 names for South American trees, including valid species and infra-specific names, synonyms, orthographical variants and common typos. The search for records based on this input list resulted in a total of 3.11 million records from 543 collections found in *species*Link, JABOT, SNDB and GBIF (Appendix A).

#### Duplicate specimen search

We standardized the notation of the collector and identifier fields. As far as possible, we assigned a unique name, following the notation of the Biodiversity Information Standards (https://www.tdwg.org), for all possible variants of notations and formats of these fields. We also standardized most of the notation of collector's number and identification year across herbaria. The search for duplicated specimen between herbaria was carried by creating codes for each specimen using the following fields: family, species, collector's last name and number, county and year of collection. Because of the great variation in collector's name notation and in the completeness of information on the collection date and locality across sources, we used four different combinations to search for duplicates: family, last name, number and county; family, number, county and year; species, last name, number and year; and last name, number, county and year (e.g., ‘Myrtaceae_Hatschbach_605_Curitiba’, ‘Myrtaceae_605_Curitiba_1947’, ‘Myrceugenia euosma_Hatschbach_605_1947’ and ‘Hatschbach_605_Curitiba_1947’). These combinations make virtually unique specimen identity and one generally retrieves a duplicated specimen if the other fails to do so.

Note that this is a quite conservative way of assigning duplicates among herbaria, because it takes into account the collection locality and the specimen taxonomy as well as the more traditional fields of duplicate search (i.e., collector’s name and number). But since we later performed a cross-validation of the available information within duplicated specimens, this conservative approach avoids problems related to false or problematic duplicate retrieval (e.g., specimens collected by two different collectors with the same last name, same series number and same county). Therefore, the ratio of duplicates found in this study should be regarded as a lower bound of the true ratio of duplicates among herbaria. In addition, if a given specimen did not have the information to construct at least one of these combinations, then the duplicate check could not be performed for this specimen, even if it would be duplicated among herbaria. Moreover, the duplicate search may not always work due to the presence of typos in the collectors' last name (e.g., Hatshcbach instead of Hatschbach), variants in its notation (e.g., Mello Barreto vs. Mello-Barreto) or due to typos, differences in notation or missing information in the collection county and/or year across herbaria. Finally, absent, anonymous or unknow collector name and/or numbers (i.e., Sellow, F., s.n. or Anonymous, s.n.) were not considered here in the duplicate search among herbaria.

### Validation of geographical coordinates and specimen locality

We then retrieved specimens with problematic geographical coordinates (i.e., zero, inverted, switched or inaccurate coordinates). We validated coordinates at country level for all countries, at state/province level for Latin America and at county levels for Argentina, Paraguay and Brazil using administrative maps available online (https://gadm.org). For occurrences with coordinates falling outside the county of reference, we calculated the distance between the coordinate and the central coordinate of the county. If this distance was up to 20 km, we kept the original coordinates, except when coordinates felt in water bodies, which were replaced by the central coordinate of the county. The 20 km threshold was set empirically as the 75% quantile of the distances between coordinates falling inside the correct county and its central coordinate. If this distance was between 20 and 40 km, we validated the occurrence if both the reference county and the given coordinates both corresponded to counties with 90% or more of its area inside or outside the Atlantic Forest limits at the scale 1:5,000,000 (IBGE, 2012; Olson & Dinerstein, 2002). For occurrences with missing coordinates but with complete information on country, state/province and county, we obtained the central coordinate of the lowest administrative level available, but only if the given county had >90% inside or outside the Atlantic Forest. The same reasoning was used to validate occurrences without coordinates, but collected in states/provinces or countries with 100% of their area completely inside (e.g., Canindeyú or Rio de Janeiro) or outside the Atlantic Forest (e.g., Amazonas or Peru). Note that for this geographical validation, we assumed that the location provided as being more precise than the coordinates themselves and that coordinates were not necessary for the geographical validation in some cases.

### Confidence levels of specimen identifications

We finally assessed the taxonomic confidence level of each specimen based on the identifier name of each specimen. Identifications were flagged as 'validated' for three different cases: (i) type specimens (e.g., isotypes, holotypes, etc), (ii) if the identification was made by a specialist of the corresponding family or (iii) if the collection was made by a specialist of the corresponding family in the case of an empty identifier field. This validation was based on a dictionary of taxonomists names per botanical family built based on information from the Harvard University Herbaria (http://kiki.huh.harvard.edu/databases), the Brazilian Herbaria Network (http://www.botanica.org.br/rbh) and the American Society of Plant Taxonomists (https://members.aspt.net). The dictionary was complemented based on our personal knowledge and internet searches and it included common variants of taxonomists names (e.g., missing initials, typos, married or maiden names - Appendix B). About two thirds of all specimens remained as 'not validated' taxonomically, so we also flagged as 'probably validated' the specimens (i) belonging to 51 taxa with low taxonomic complexity (i.e., easy to identify, such as the Atlantic Forest trees *Araucaria angustifolia* or *Piptadenia gonoacantha* - Appendix C) and (ii) identification performed by plant taxonomists in general. We assume that if a taxonomist identified a specimen from a family outside his specialties, he is confident in the identification, and thus the identification is more reliable than the ones made by non-taxonomists. Because the inclusion of 'probably validated' specimens may increase identification errors, these specimens were not considered in constructing the Atlantic Forest checklist.

#### Cross-validation within duplicated specimens

For groups of duplicates in which we had a high degree of certainty in their grouping (e.g., all four different codes of duplicate search were equal), we combined the information from the different occurrences in order to obtain the information with the best quality possible. We first combined the information on the geographical validation and the occurrence position in respect to the Atlantic Forest limits. For instance, if one of the occurrences had its coordinates validated at county level, the coordinates of all duplicates were flagged as validated at county level. The information on the sampling locality and position in respect to the Atlantic Forest limits were also completed within the group of duplicates. For plotting purposes only, the coordinates of the records validated at the best resolution possible, were averaged to create a mean coordinate for the group of duplicates. In the case of records with coordinates validated at county level but with contrasting positions in respect to the Atlantic Forest limits (e.g., inside and outside), the group of duplicates was not used for further analysis.

We also combined information on the species identification and taxonomic confidence level from duplicated specimens. For instance, if one of the duplicates had a species identification flagged as 'validated', the species identification of all duplicates was then flagged as 'validated'. In the case of different identifications within a same group of duplicates, the identification flagged as validated was taken as the valid one. In the case of conflicting identifications between specialists in the same family, the record with the most recent identification year was chosen as the valid one for a given group of duplicates.

#### Detection of spatial outliers in species occurrences

Although we have performed an in-depth validation and cross-validation of the geographical coordinates, some occurrences too distant from the species core distributions can remain. One common cause to these spatial outliers is related to collections that were properly identified by a family specialist but that were not declared as being taken from cultivated individual in the specimen notes. This was often the case for samples taken from individuals inside botanical gardens. We could exclude all records collected from botanical gardens, but we decided not to do so because many botanical gardens in South America have both cultivated trees (i.e., arboretum) and natural forest fragments.

We used an approach based on spatial outliers combining to different methods to detect outliers (Liu et al., 2018): the classic and the robust Mahalanobis distances. The two methods take into account the geographical centre of the species distribution and the covariance between the occurrences. But they vary in the way the covariance matrix of the distribution is defined: the classic distance uses an approach based on Pearson's method, while the robust distance uses a Minimum Covariance Determinant estimator. Next, based on the empirical distribution of some

Atlantic Forest species with very different number of occurrences and spatial distribution patterns, we observed that occurrences truly outside the species ranges often had classic and robust Mahalanobis distances above 3 and 16. We used these thresholds to remove occurrences for all species (about 49,000 or 0.8% of occurrences excluded), which for most species resulted in no exclusion at all (very conservative outlier removal). As expected, species cultivated by man had more spatial outliers than other species. So, for a list of 42 cultivated species (Appendix C), we used more restrictive thresholds of 2.5 and 12, respectively. These thresholds still remain very conservative (exclusion of about 2900 or 1.4% of the occurrences for cultivated species). Therefore, some outliers may have remained in the dataset.

### Descriptive results of the validation process of the occurrence data

From the total of 3.11 million records retrieved for the list of names for South American trees, 92.2% had their geographical location validated at least at country level (63.8% at county or locality levels; 13.7% at state or province level; 14.7% at country level only). About 5% of the occurrences had no coordinates at all, and for the remaining 3% we could not perform the validation (e.g., no locality information at all). Regarding the confidence levels of the identifications, 40.3% of the occurrences had their species identification validated by taxonomic experts (Appendix B). In addition, we found that about 24% of all records were duplicated between two or more collections. Thus, we ended with a total 735,128 non-duplicated, geographically and taxonomically validated occurrences. From this total, 544,252 occurrences could be unambiguously related to specimens collected inside or outside the Atlantic Forest limits. Occurrences in the transitions of the Atlantic Forest to other South American domains represented 49,668 additional occurrences. All other tree names that were found for Atlantic Forest but were not included in the checklist are given in Appendices D and E, along with the reason of their exclusion.

One main issue for the data validation process was the lack of information on collection locality at county/municipality levels (31% of the records). These fields are essential to validate the original coordinates at finer spatial scales or to retrieve missing coordinates from gazetteers. Updating or completing these fields in the original herbarium labels should be straightforward and would avoid major data leakage (*sensu* Townsend Peterson et al. 2018). Missing information on the identifier name (28%) had a similar impact on the taxonomic validation as occurrences not identified by family specialists (32%). Mitigating this source of data leakage, however, would require much more human and economic resources when compared to completing locality information. Here, we had an increase of 23% in the number of valid occurrences by considering specimens with both ‘validated’ and ‘probably validated’ taxonomy (see ‘Confidence levels of specimen identifications’ above), leading to a combined total of 949,721 records (Appendix C).

### General description of the Atlantic Forest tree flora

As described in the main text, we found 252,911 valid records inside the Atlantic Forest, which revealed a total of 5044 arborescent species, including 4054 tree species (Appendix C). The ten most species-rich families for the Atlantic Forest arborescent flora were Myrtaceae (679 species), Fabaceae (655), Rubiaceae (327), Melastomataceae (289), Lauraceae (220), Euphorbiaceae (174), Asteraceae (167), Solanaceae (161), Annonaceae (117) and Malvaceae

(113). These families represented 58% of all the arborescent species recorded for the Atlantic Forest. Regarding only tree species, the top 10 families remained relatively the same (Myrtaceae: 626; Fabaceae: 544; Rubiaceae: 221; Lauraceae: 217; Melastomataceae: 211; Euphorbiaceae: 126; Annonaceae: 106; Sapotaceae: 91; Malvaceae: 89; and Sapindaceae: 78). Among families with more than 25 species in the Atlantic Forest arborescent flora, some had high average endemism level (>70%), namely Monimiaceae (95%), Symplocaceae (85%), Poaceae: Bambusoideae (84%), Araliaceae (83%), Myrtaceae (81%), Cyatheaceae (80%), Proteaceae (79%), Aquifoliaceae (78%), Lauraceae (76%), Rutaceae (74%), Melastomataceae (72%) and Annonaceae (71%).

We found much less valid occurrences for the Atlantic Forest parts in Argentina and Paraguay (2367 and 4049, respectively) than in Brazil (246,495 occurrences). This was expected from the smaller area of Atlantic Forest in these countries. But, most of the Atlantic Forest areas in these two countries have low sampling coverage (Figure 3a), with many Atlantic Forest grid cells with no occurrences at all in these countries. In addition, the ratio of validated vs. retrieved records for Argentina (21%) was also smaller than for Brazil and Paraguay (38 and 39%, respectively). Therefore, the number of species found for Argentina (303) and Paraguay (462) are most probably underestimated. For the Brazilian part of the Atlantic Forest, we retrieved a total of 5025 species.

**Notes on endemism level calculation and species classification**

We calculated the endemism level using (i) only taxonomically ‘validated’ records (Figure S4) and (ii) both taxonomically ‘validated’ and ‘probably validated’ for each species occurring in the Atlantic Forest (Figure 1, Appendix F). These endemism levels and thus the classification of species must be used with some caution for different reasons. First, different life forms and regions are often not collected with the same intensity (ter Steege et al., 2011; Schmidt-Lebuhn et al., 2013; Osazuwa-Peters et al., 2018). For instance, *Euterpe edulis* Mart. (palm-heart) is possibly the most abundant arborescent species in the Atlantic Forest, being relatively easy to found fertile and to collect in the field. Nonetheless, it is ranked 1682 out of 4989 in terms of records available to compute its endemism level (about 130 records). One reason for this discrepancy between the abundance of the species in the field and the number of herbarium records available is the reluctance to collect and herborize palm leaves, fruits and flowers, an issue which is also true for other life forms such as cacti, tree ferns, and woody bamboos. In addition, *E. edulis* is relatively easy to identify in the field and it is the only representative of the genus in the Atlantic Forest, so it is often not collected. Moreover, there may be a collection bias outside its core distribution, where researchers may be more prone to collect the species to confirm its identification in the herbarium. Therefore, this species is classified here as a widespread species, although it is most likely a near endemic.

Second, some emblematic and/or ornamental species may have been collected from cultivated individuals in botanical gardens, city squares or house gardens. If the collector did not declare as such or was unaware that the individual was cultivated, then the record could not be detected and removed and it entered the computation of the endemism level. As described above, the detection of spatial outliers should decrease these types of problems, but it does not remove all. The inclusion of these cultivated records could lead to an underestimate of the species endemism level. This was probably the case of *Araucaria angustifolia* (Bertol.) Kuntze and

*Paubrasilia echinata* (Lam.) Gagnon, H.C. Lima & G.P. Lewis, two of the most emblematic Atlantic Forest endemics, which are often cultivated outside their natural range.

Moreover, some of the mismatches between the endemism classification proposed here and the classifications found in the Brazilian Flora 2020 project comes from species typically associated to specific "habitat islands" within the Atlantic Forest hotspot. This is the cases for species, such as *Cinnamomum erythropus* (Nees & Mart.) Kosterm., *Micropholis emarginata* T.D. Penn., *Piptolepis monticola* Loeuille, *Swartzia bahiensis* R.S. Cowan and *Vantanea morii* Cuatrec., which are mainly found in rocky highland fields, areas which are often associated with the Cerrado domain. Therefore, one should bear in mind that the classification proposed here results from a calculation including records in all habitats and vegetation types within the Atlantic Forest limits (IBGE, 2012; Olson & Dinerstein, 2002).

Finally, the thresholds presented in the main text are the most conservative breaking point obtained from a piecewise regression analysis. For near endemism, a second, more inclusive optimum threshold was found at 85.5% of endemism level. The same threshold using only taxonomically 'validated' records was also slightly smaller (88%; Figure S4). For occasional species, a second threshold was found at 15.3%. The reason for these differences is because the curves of mismatches became almost flat towards their lowest parts, meaning that most of the values within this flat part could be equally accurate. We tried to solve this limitation of the piecewise regression by finding the value of endemism level where the first derivate of the curves is zero. But since the derivate values were too variable in respect to the 1% threshold values in these flatter parts of the curves, assigning a single and accurate first derivate proved to be challenging as well. Therefore, the thresholds of endemism level presented in the main text should be seen as safe, conservative thresholds. So, a more practical near-endemic threshold is somewhere between 86 and 90%, most probably at 88%, which would change the overall endemism ratio of the Atlantic Forest tree flora from 45 to 47%.

### Performance of the diversity metrics per grid-cell

We obtained for each 50×50 km grid cell four metrics: observed species richness (S), corrected weighted endemism (WE), rarefied/extrapolated richness ($S_{RE}$) and the effective richness ($S_E$). WE and $S_{RE}$ are described in the main text. The metric S is the simple count of species per cell. $S_E$ is the asymptotic estimate of the Simpson's index, which downweighs rare species, typically over-represented in herbarium data (ter Steege etal., 2011). Both $S_{RE}$ and $S_E$ were calculated based on Hill numbers (q=0 for $S_{RE}$ and q=2 for $S_E$) and frequencies of species occurrences per cell (Chao et al., 2014). All of these diversity metrics have limitations, because herbarium data is typically aggregated in space and it often represent a non-random sample of the local plant communities (Osazuwa-Peters et al., 2018; Schmidt-Lebuhn et al., 2013; ter Steege et al., 2011). However, these metrics are indicated to control for sample size while assessing spatial patterns of biodiversity (Engemann et al., 2015; Osazuwa-Peters et al., 2018).

We evaluated the relationship of diversity metrics with sampling intensity (i.e., number of occurrences per cell). As describe in the main text, these evaluations were performed using spatial regression models (i.e., linear regression with spatially correlated errors), using an exponential variogram model. As expected, the number of endemic and occasional species per grid cell were highly correlated with the sampling intensity (Figure S2). Other diversity measures were also highly correlated to sampling intensity (Figure S3), even if we consider only the grid cells with sampling coverage above 80% (results not shown). The metrics WE and $S_{RE}$ were the less correlated with the sampling intensity (Figure S3). We thus selected $S_{RE}$, which is given in a unity more straightforward to understand (i.e., number of species), as the metric to delimit the centres of endemism and occasional species in the Atlantic Forest, as shown in the main text (Figures 2 and 3).

### Delimiting centres of species diversity

The delimitation of the centres of diversity was performed using ordinary kriging on log-transformed diversity metrics. We used an exponential variogram model, nugget effect and we set a maximum of 20 nearest observations to perform the interpolation at the centres of a 5×5 km grid. Spatial regressions (described above) and kriging analysis included only the grid cells meeting the following criteria: overall sample coverage between the 25–50% quantiles of the distribution of all cells and ≥100 occurrences, sample coverage between the 50–75% quantiles and ≥50 occurrences; or sample coverage ≥75% quantile and ≥30 occurrences. About 48% of the total of 2695 50×50 km grid cells met these criteria. Consequently, analyses included grid cells with a median of 254 occurrences (range: 37–10,744) and 70.9% of sampling coverage (range: 34.6–98.2%). Only 11% of the grid cells included in the analysis had less than 100 occurrences. For each group of species, we present the centres of diversity based on observed species richness (Figure S6), corrected weighted endemism (Figure S7) rarefied/extrapolated species richness (Figure 3) and effective species richness (Hill's q= 2; Figure S8).

We found that the diversity of pure endemics in the Atlantic Forest was concentrated in the rainforests along the Brazilian coast, between Santa Catarina and Bahia states. Most of metrics showed some discontinuity in the centres of endemism along the coast (Figure 3 and Figures S6-S8), corresponding to the regions of north of Rio de Janeiro and Espírito Santo states, corresponding to areas with higher rainfall seasonality and with the occurrence of lowland seasonal forests (Rodal et al., 2006). The influence of more seasonal climates and enclaves of seasonally dry forests may also be the reason why we found no clear centre of endemic diversity in northeast Brazil above the north of Bahia state.

The corrected weighted endemism (WE), which takes into account the range of species occurring in the grid cells, was the metric that suggested the most different delimitation of centres of endemism (Figure S8). This metric suggested that the ES-BA centre of endemism should be wider, while the SP-RJ/PR-SC centres of endemism should be narrower or non-existent. This means that the ES-BA centre of endemism has more endemics with narrow ranges (i.e., highly restricted endemics), while the SP-RJ/PR-SC centres of endemism has more endemics with wider ranges. The high diversity of occasional species in the central part of the Atlantic Forest, suggested by this metric (Figure S8d) is probably due to the fact that species ranges were obtained only within the extent of our grid, which does not cover most of the Amazon. So, species with disjunct distributions between the Amazon and the Atlantic Forest are overestimating the metric estimation.

Although collections may actually be more abundant in more diverse areas (i.e., collection bias in areas known to be richer in species), the strong correlation found between diversity measures and sampling intensity suggests that the number of occurrences available for some parts of the Atlantic Forest is still very limited to rule out a possible sampling bias (Murray-Smith et al., 2009). These areas include the Atlantic Forest parts in Argentina, Paraguay and in the Brazilian states of Mato Grosso do Sul, Goiás, Piauí and Ceará, as well as the central and western parts of Rio Grande do Sul, Santa Catarina, Paraná, São Paulo and Bahia (Figure 3a). This means that some areas of the Atlantic Forest could be included as centres of endemism and some of the discontinuities could disappear with the accumulation of new collections. Therefore, the proposal of centres of endemism should be interpreted in the light of the occurrence data currently available (Murray-Smith et al., 2009).

**Software and packages used for data analysis**

All codes and analysis in this study were performed using R (R Core Team, 2018). We used the contributed packages 'RCurl' for data download (Lang & R Core Team, 2018), 'finch' for Darwin core parsing (Chamberlain, 2018), 'flora' and 'Taxonstand' for species name checking (Carvalho, 2017; Cayuela, Stein, & Oksanen, 2017), 'mvoutlier' for spatial outlier detection (Filzmoser & Gschwandtner, 2018), 'segmented' for piecewise regression (Muggeo, 2008), 'iNEXT' for diversity metrics (Hsieh, Ma, & Chao, 2016), 'nlme' and 'gstat' for spatial regressions and kriging (Gräler, Pebesma, & Heuvelink, 2016; Pinheiro, Bates, DebRoy, Sarka, & R Core Team, 2018).

### SUPPLEMENTARY FIGURES

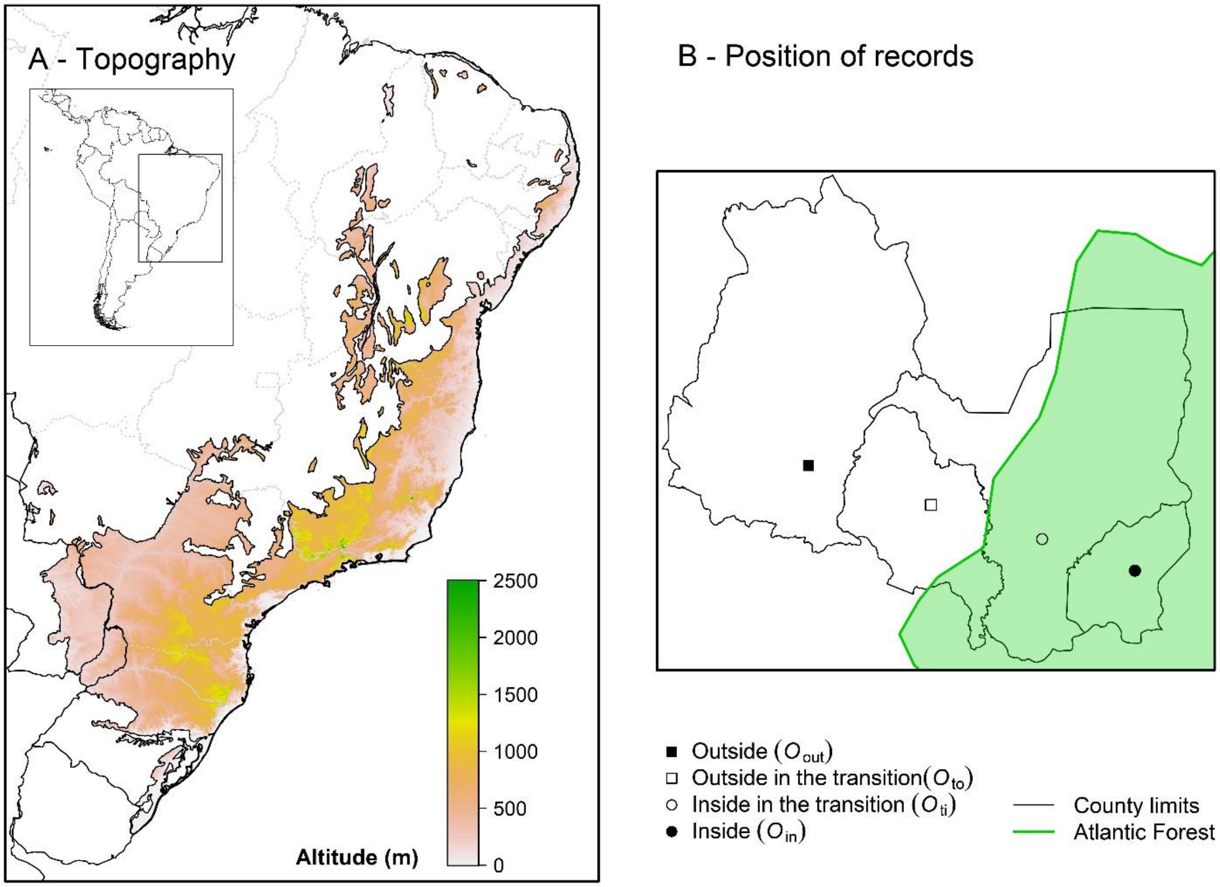


**Figure S1**. The (A) location and topography of the Atlantic Forest in Eastern South America and (B) the four types of possible position of records in respect to the Atlantic Forest limits. Each record was assigned as being inside ($O_{in}$), outside ($O_{out}$) or in the transition of the Atlantic Forest to other domains ($O_{ti}$ or $O_{to}$) and these types of position were used to calculate the species level of endemism. In panel A, black and grey lines mark the limits of South American countries and of Brazilian states, respectively. In panel B, the shaded green area represents the Atlantic Forest, while the rest of the map (white area) represents a different domain (e.g., *Cerrado* or *Caatinga*).

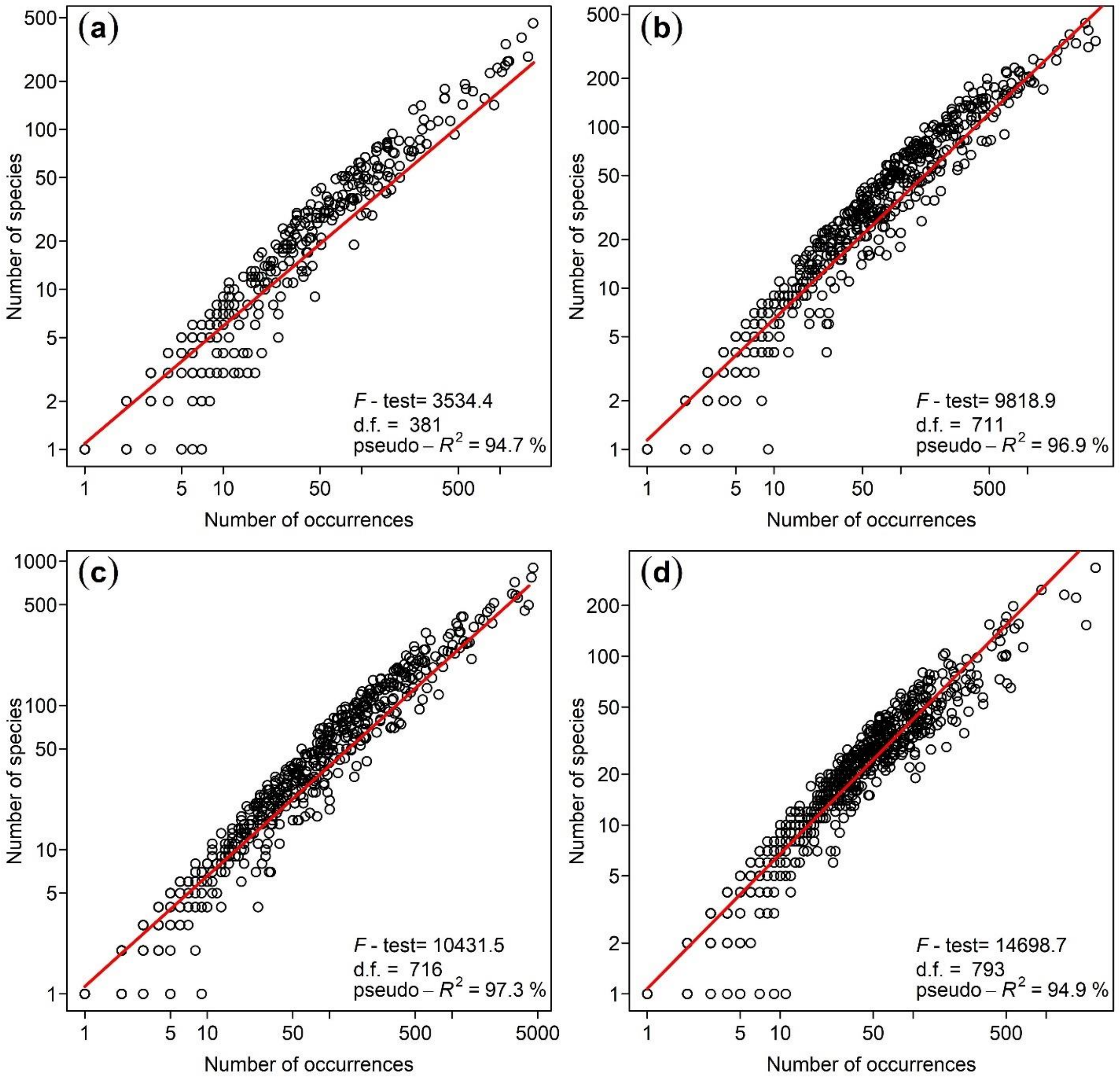


**Figure S2**. Relationship between the number of species and the number of occurrences per grid cell for (a) pure endemics, (b) near endemics, (c) pure + near endemics and (d) occasional species. For each group of species, we present the mean prediction of the spatial regression model (red lines) and the summary statistics of each spatial regression model (bottom right; d.f.= degrees of freedom). Although the scatterplot present values for all grid cells, the spatial regression analysis was performed only for grid cells meeting some minimum criteria of sampling coverage (see Supplementary Methods). Both axes are in logarithmic scale.

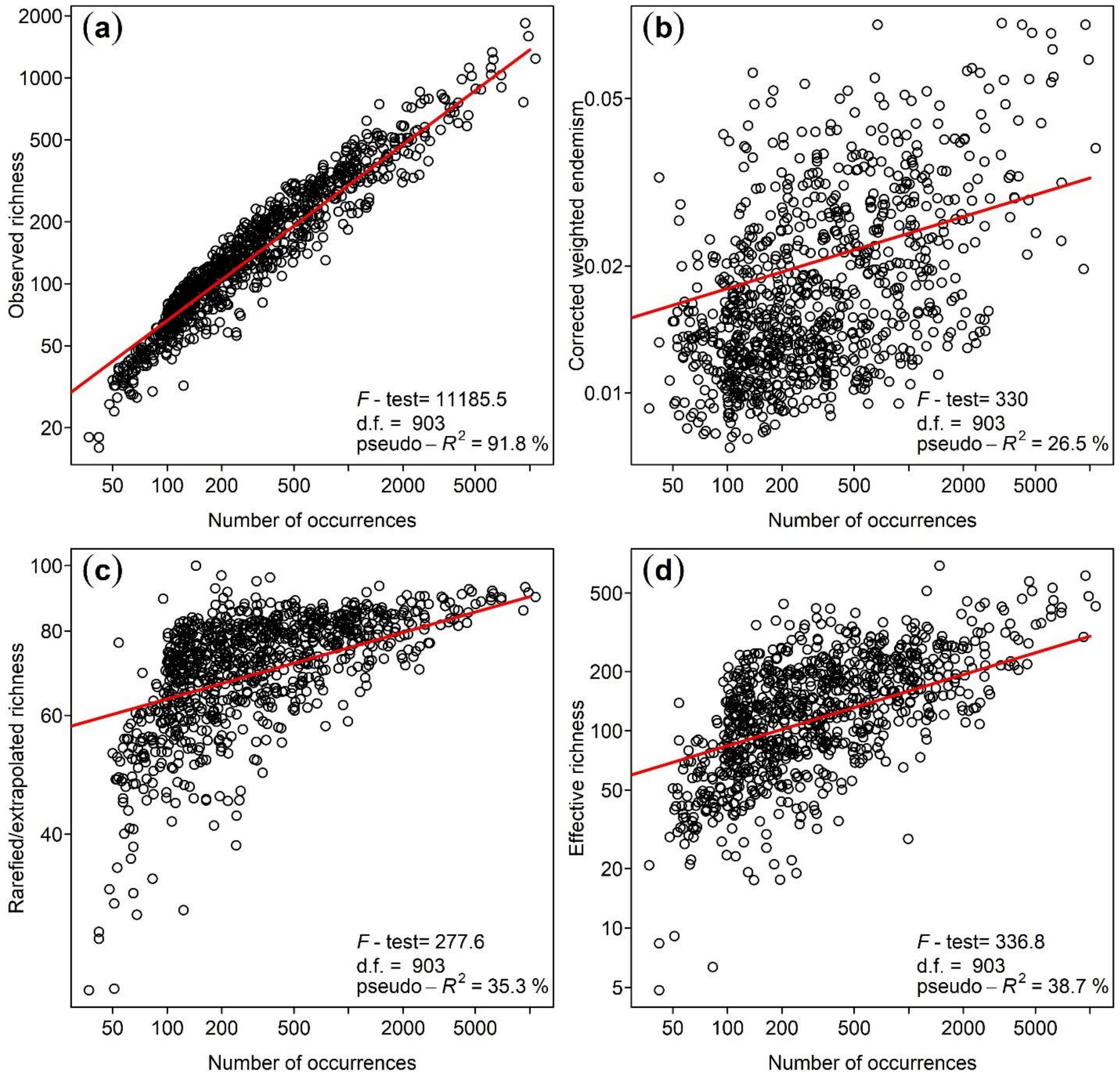


**Figure S3**. Relationship between the number of occurrences per grid cell and the four measures of diversity evaluated here, namely the (a) observed species richness, (b) corrected weighted endemism, (c) rarefied/extrapolated species richness and (d) effective species richness (Hill's q= 2). For each diversity metric, we present the mean prediction of the spatial regression model (red lines) and the summary statistics of each spatial regression model (bottom right; d.f.= degrees of freedom). Although the scatterplot present values for all grid cells, the spatial regression analysis was performed only for grid cells meeting some minimum criteria of sampling coverage (see Supplementary Methods). Both axes are in logarithmic scale.

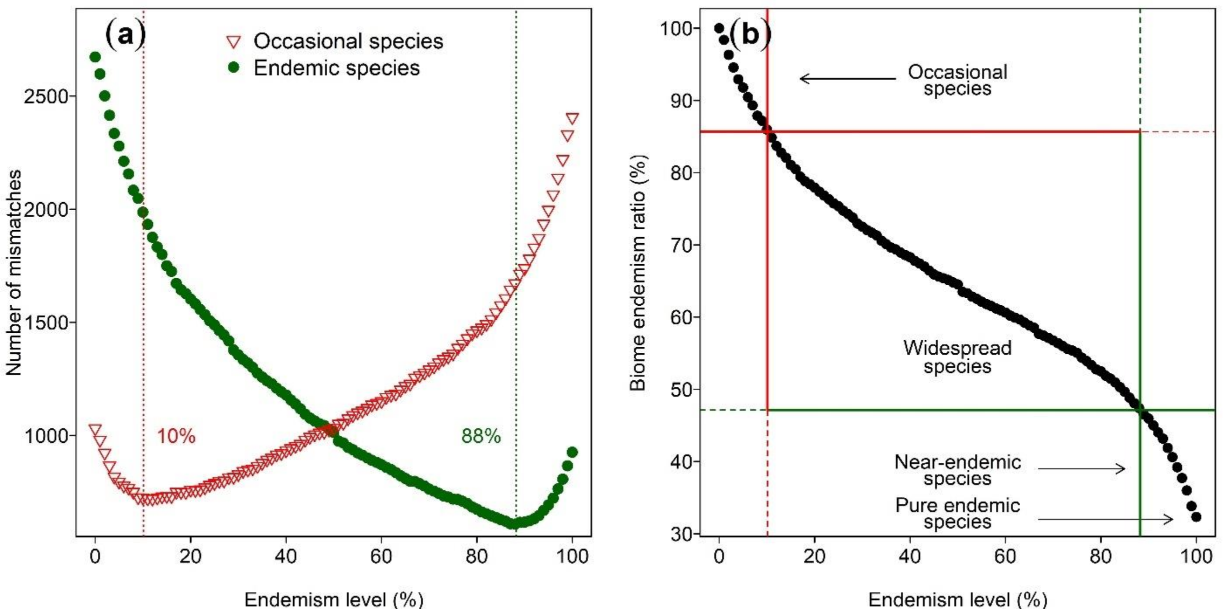


**Figure S4**. Defining near endemic and occasional species using only herbarium records with taxonomy flagged as 'validated' for the Atlantic Forest. For both endemic (green circles) and occasional species (red triangles), we present (a) the optimum endemism levels (vertical dashed lines) from the distribution of mismatches between the observed and the Brazilian Flora 2020 classifications and (b) the overall endemism ratio of the Atlantic Forest in intervals of 1% (*x*-axis in both panels). The results of this figure were the result of the same analysis that generated Figure 1 in the main text, but using a different subset of valid occurrences.

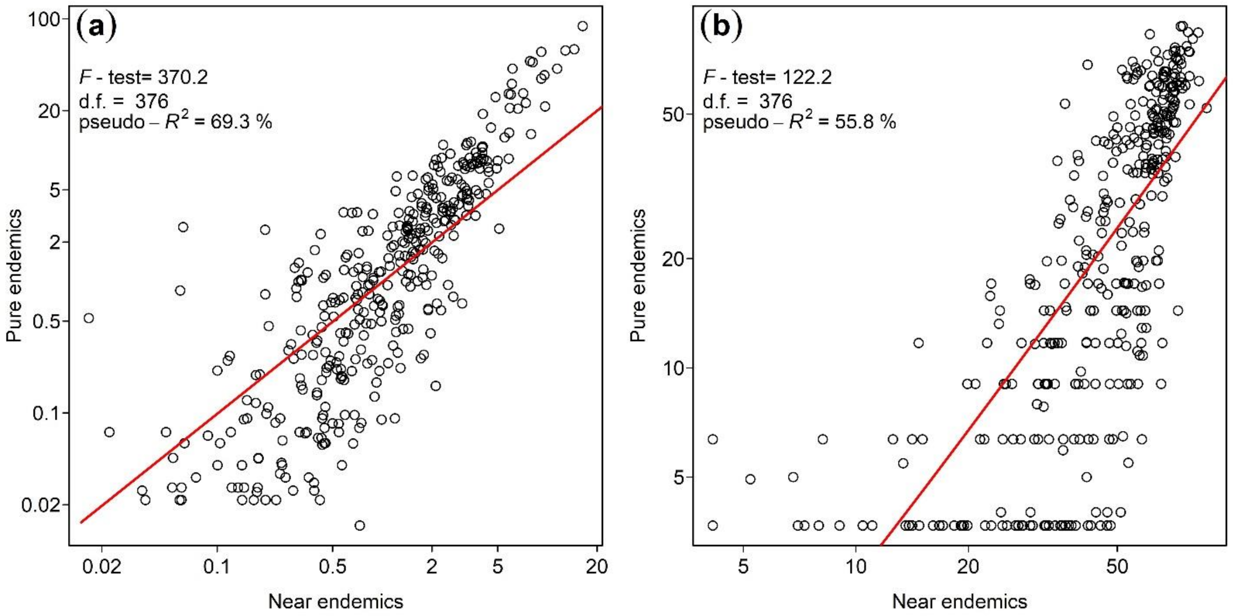


**Figure S5**. Relationship between the number of near and pure endemic species per grid cell regarding the two best diversity metrics: (a) corrected weighted endemism and (b) rarefied/extrapolated species richness for 100 records. For both metrics, we present the mean prediction of the spatial regression model (red lines) and the summary statistics of each model (top left; d.f.= degrees of freedom). Although the scatterplot present values for all grid cells, the spatial regression analysis was performed only for grid cells meeting some minimum criteria of sampling coverage (see Supplementary Methods). Both axes are in logarithmic scale.

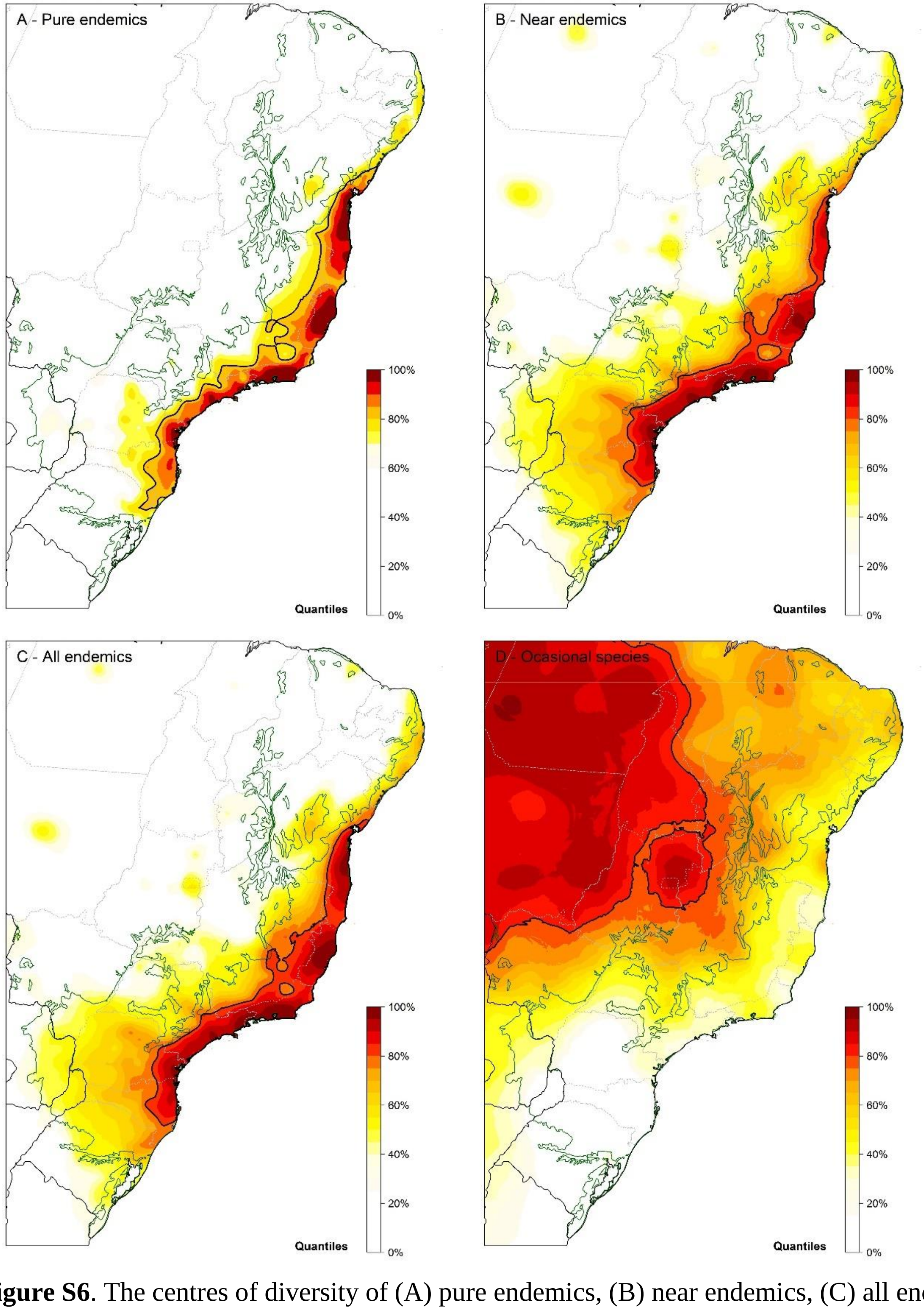


**Figure S6**. The centres of diversity of (A) pure endemics, (B) near endemics, (C) all endemics (pure + near) and (D) occasional species in the Atlantic Forest (green line), based on the observed species richness. The colour scale represents the 5% quantiles of the metric distribution, from 0% (white) to 95-100% (dark-red). Black and grey lines mark the limits of South American countries and of Brazilian states, respectively.

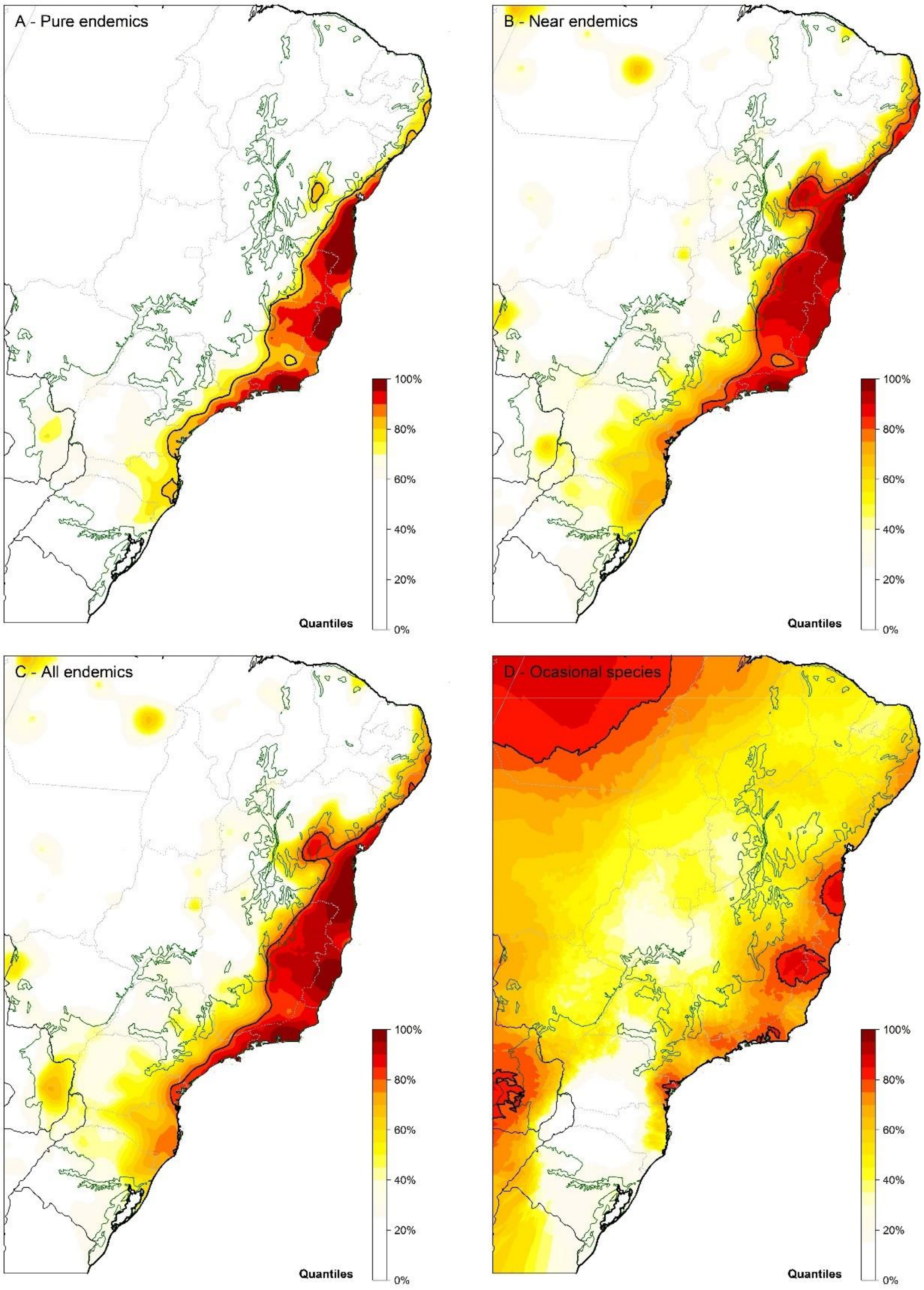


**Figure S7**. The centres of diversity of (A) pure endemics, (B) near endemics, (C) all endemics (pure + near) and (D) occasional species in the Atlantic Forest (green line), based on the corrected weighted endemism. The colour scale represents the 5% quantiles of the metric distribution, from 0% (white) to 95-100% (dark-red). Black and grey lines mark the limits of South American countries and of Brazilian states, respectively.

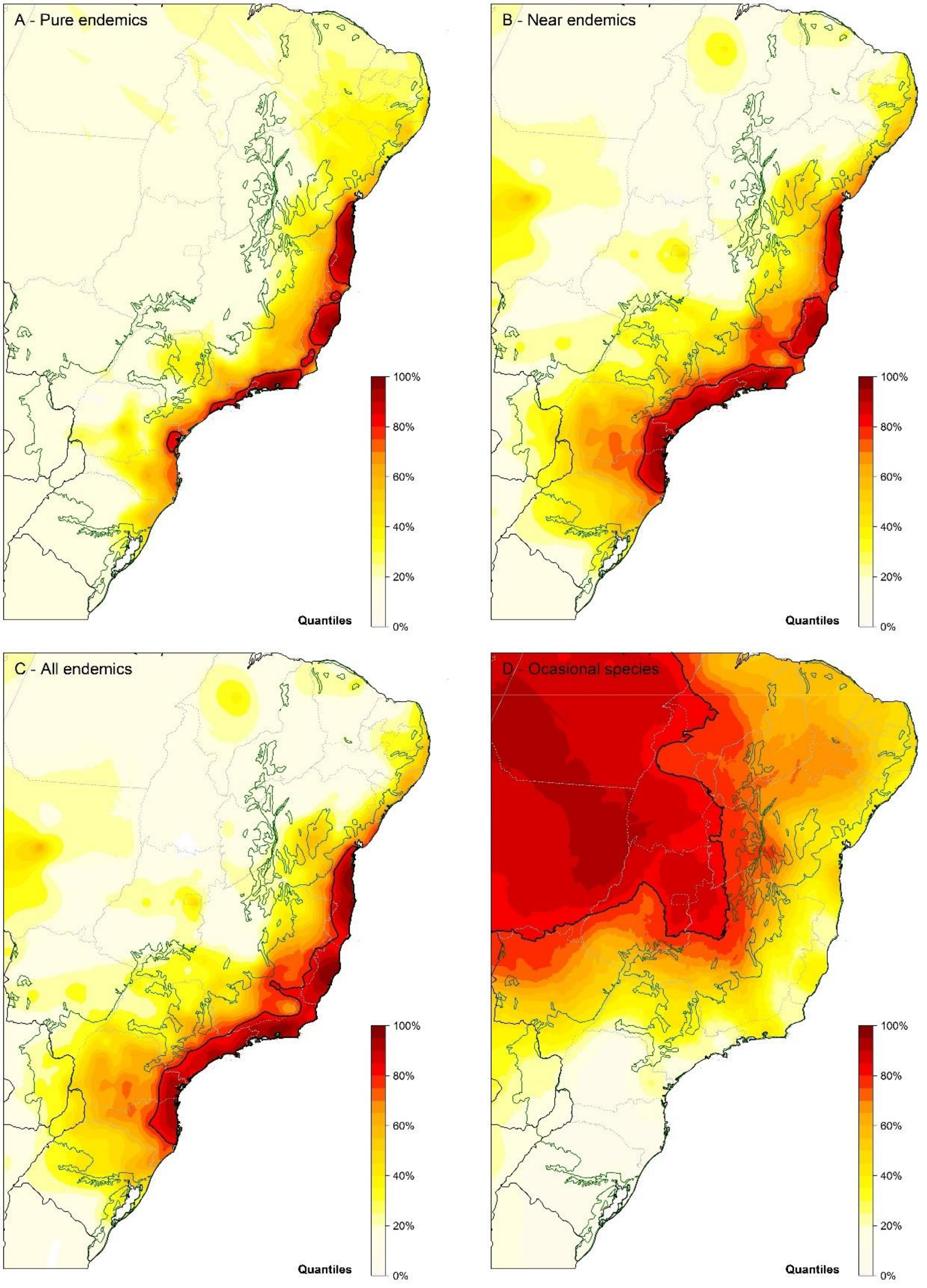


**Figure S8**. The centres of diversity of (A) pure endemics, (B) near endemics, (C) all endemics (pure + near) and (D) occasional species in the Atlantic Forest (green line), based on effective species richness (Hill's q= 2). The colour scale represents the 5% quantiles of the metrics distribution, from 0-5% (light-yellow) to 95-100% (dark-red). Black and grey lines mark the limits of South American countries and of Brazilian states, respectively.
